# Interaction of the Immune System TIM-3 Protein with a Model Cellular Membrane Containing Phosphatidyl-Serine Lipids

**DOI:** 10.1101/2023.12.24.573254

**Authors:** Anastasiia Delova, Andreea Pasc, Antonio Monari

## Abstract

T cell transmembrane, Immunoglobulin, and Mucin (TIM) are important immune system proteins which are especially present in T-cells and regulated the immune system by sensing cell engulfment and apoptotic processes. Their role is exerted by the capacity to detect the presence of phosphatidyl-serine lipid polar head in the outer leaflet of cellular membranes (correlated with apoptosis). In this contribution by using equilibrium and enhanced sampling molecular dynamics simulation we unravel the molecular bases and the thermodynamics of TIM, and in particular TIM-3, interaction with phosphatidyl serine in a lipid bilayer. Since TIM-3 deregulation is an important factor of pro-oncogenic tumor micro-environment understanding its functioning at a molecular level may pave the way to the development of original immunotherapeutic approaches.

## Introduction

The family of T cell transmembrane, Immunoglobulin, and Mucin (TIM) proteins plays a pivotal role in the intricate regulation of both innate and adaptive immunity.^[1,2]^ Among the different members, TIM-1, TIM-2, TIM-3, and TIM-4 have been particularly considered.^[3–5]^ Initially TIM-1 and TIM-4 have been recognized as receptors for phosphatidylserine (PtdSer). Yet, the same interaction pattern, and the high affinity towards PtdSer have been recently evidenced for TIM-3.^[6,7]^ Indeed, while in normal cells PdtSer bearing lipids are mainly localized on the inner membrane leaflet, they becomes exposed on the outer leaflet following cell activation or apoptotic signals,^[8,9]^ Therefore, they represent a signal of cell engulfment and activate the immune response through their interaction with TIM proteins.^[10–13]^ Yet, the binding of PtdSer to the IgV domain of TIM not only favors the recognition of apoptotic cells but has also wider implications for the global immune response,^[14,15]^ for instance by triggering potent anti-inflammatory effects,^[16–19]^ which are essential for maintaining immune tolerance. In contrast with the other members of the family, TIM-2 does not bind to PtdSer and, thus, serves a different function, mainly related to T-helper type 2 response and autoimmunity, as well as iron blood level.^[20–23]^

TIM-3 is preferentially expressed on T-helper 1 (Th1) and T-cells 1 (T-cells) but is also found on dendritic cells,^[24]^ and is recognized as a marker of maturation in natural killer (Nk) cells.^[25–29]^ The interaction of TIM-3 with its ligand usually generates an inhibitory signal culminating in the apoptosis of Th1 cells and the production of cytokine.^[30,31]^ Furthermore, TIM-3 is usually co-expressed with other check point modulators also leading to the T-cells exhaustion in tumors.^[32]^ Indeed, TIM-3 may lead to the functionality loss of lymphocytes in the tumor microenvironment,^[33–35]^ which correlates with a bad prognosis and, therefore, is recognized as an important cancer biomarker.^[36–39]^ Globally, all these observations underscore the potential of TIM-3 in modulating immune responses, particularly in the context of autoimmunity, but also the fact it represents a potential target in cancer immunotherapy.^[40]^ Indeed, in vivo and pre-clinical results supports the role of the blockade of TIM-3 in promoting tumor immunity, even in advanced cancers.^[41–43]^

From a structural point of view,^[44,45]^ as schematized in Figure 1, TIM-family members comprise rather large transmembrane proteins, which protrude from the lymphocyte’s membrane, and are composed of several domains. As shown in Figure 1, TIM-3, and more generally TIM proteins, are composed of a disordered cytosolic domain, a transmembrane segment, anchoring the protein to the immune system cell, and the hydrophilic mucin domain, which also presents glycosylation sites.^[46]^

**Figure 1.**
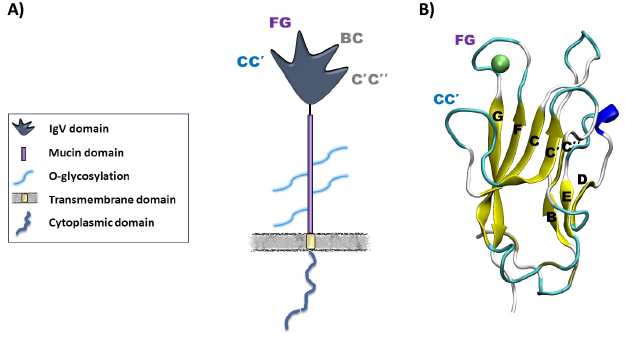
A) Schematic representation of the different domains of the TIM-3 transmembrane protein. B) Structure of the Ig-domain of TIM-3 in cartoon representation. The Ca^2+^ ion in the PtdSer binding pocket is also evidenced in can der Walls representation.

Finally, the N-terminal immunoglobulin-like (IgV) domain, comprising the PtdSer binding site, is tethered to the mucin domain and senses the outer leaflet of the target cells. The globular IgV-domain (Figure 1) presents four flexible and disordered loops, named CC’, FG, BC, and C’C’’. While the PdtSer binding pocket, is surrounded by the two rather hydrophobic CC’ and FG loops, which favors the interaction with the target cell lipid membrane, the core of the IgV domains, is composed of two antiparallel β-sheets sub-domains. As can be appreciated in Figure 1B, the two β-sheets sub-domains are asymmetric and the shorter BED faces the longer AGFCC’C’’ sub-domain, which also slightly encompasses the binding pocket. Both structural units clearly confer a structural stability and rigidity to the whole protein core. A further defining characteristic of the TIM-3 IgV domains is the presence of six conserved cysteine residues, involved in disulfide bridges also locking two β-sheets. The disulfide bonds are important to stabilize the extended CC’ loop, while orienting it towards the GFC motif. Indeed, in PtdSer-binding TIMs, such as TIM-1, TIM-3, and TIM-4, the CC’ loop aligns parallel to the FG loop of the IgV domain. Indeed, such structural alignment provides a pocket which allows the recognition and binding of the substrate.^[6]^

The binding of PdtSer is also mediated by the presence of a calcium ion in the binding pocket, leading to a metal ion-dependent ligand-binding site (MILIBS). The FG-CC’ loop which encompasses the binding pocket plays a pivotal role in favoring the recognition of PdtSer, since different residues of these loops are directly interacting with the lipid polar head, as it will be detailed in the following. Furthermore, the binding of PdtSer induces significant changes in the receptor’s structure. For instance, K. Weber and coworkers have shown that the binding of PtdSer to human TIM-3 induces modifications of the electrostatic environment of the FG-CC’ ligand-binding cleft.^[47]^ This change results in the formation of local salt bridges stabilizing the protein/ligand complex.^[47]^

On a more global level the binding of PdtSer, induces the phosphorylation of conserved cytosolic tyrosines, which leads to the release of the HLA-B-associated transcript 3 (BAT3) mediating the immune response.^[48]^ However, the precise mechanisms of the TIM-3-dependent cellular signaling have not been completely elucidated, yet. Interestingly, it has been recently shown that the affinity of PdtSer lipids towards TIM-3 binding may be modulated by the photoisomerization of molecular photoswitches embedded on the hydrophobic lipid tails by the absorption of suitable wavelengths.^[29]^ Such an outcome, is particularly fascinating since it can pave the way to the development of photo-immune therapeutic approaches. Yet, the full exploitation of these potentialities should necessitate the precise knowledge of the fine molecular mechanism leading to PdtSer recognition and TIM-3 activation, which remains partially elusive.

In particular, while the crystal structures of the PdtSer, i.e. the polar head, bound to TIM-3 is available,^[29]^ the interaction of the IvG domain with the lipid membrane, and with the lipid hydrophobic tails is less studied. For these reasons, in the present contribution we will resort to long-range molecular dynamic (MD) completed with enhanced sampling methodologies to unravel the factors leading to PdtSer recognition and binding. More specifically, we will assess the interaction of TIM-3 IvG domain with a model lipid bilayer, mimicking the composition of a eukaryotic cell membrane, and we will also determine the thermodynamic driving force leading to PdtSer recruitment. Furthermore, the perturbation of the IvG domain upon the binding of the ligand will also be analyzed and rationalized.

## Results and Discussion

To better understand the modification brought upon by the binding with PdtSer-containing lipids, such as POPS, on both global and local structural arrangements of the IvG domain in complex with a lipid bilayer, we simulated two systems: i) the IvG domain without any POPS lipid in the binding cleft, hereafter referred as the unbound state; ii) the IvG domain with a POPS unit in the binding pocket, hereafter styled the bound state. Details on the construction of the two systems are provided in the Computational Methodology section.

As shown in Figure 2 both the simulation for the POPS-bound and unbound state yielded a stable aggregate with the lipid bilayer. More specifically, and as expected the IvG domain behaves as a peripheral membrane protein and indeed, the CC’ and FG loop inserts into the polar head region of the bilayer. In contrast, the other two loops, i.e. BC and C’C’’, only develops some temporary interactions with the hedges of the polar head regions, being mostly solvent exposed. Interestingly, this conformation of the membrane protein aggregate is clearly optimal for sensing the presence of PdtSer lipids since it positions the binding pocket at the core of the polar head regions, as shown by the position of the Ca^2+^ ion, with respect to the closest lipids which is shown in Figure 2 as a van der Waals sphere. Interestingly, the loosely bound conformation of the IvG domain, which interacts only superficially with the lipid bilayer, is also coupled to an important mobility of the protein around the plane defined by the membrane polar heads. This aspect is also important since it allows the binding pocket to sense and probe different spatial areas of the membrane, and favors the encountering with the natural ligand. As can be also inferred from Figure 3C the hydrophobic tails of the bound POPS lipid should instead remain in significantly straight conformations, also to avoid steric clashes with the FG and CC’ loops, and more specifically with the binding cleft.

**Figure 2).**
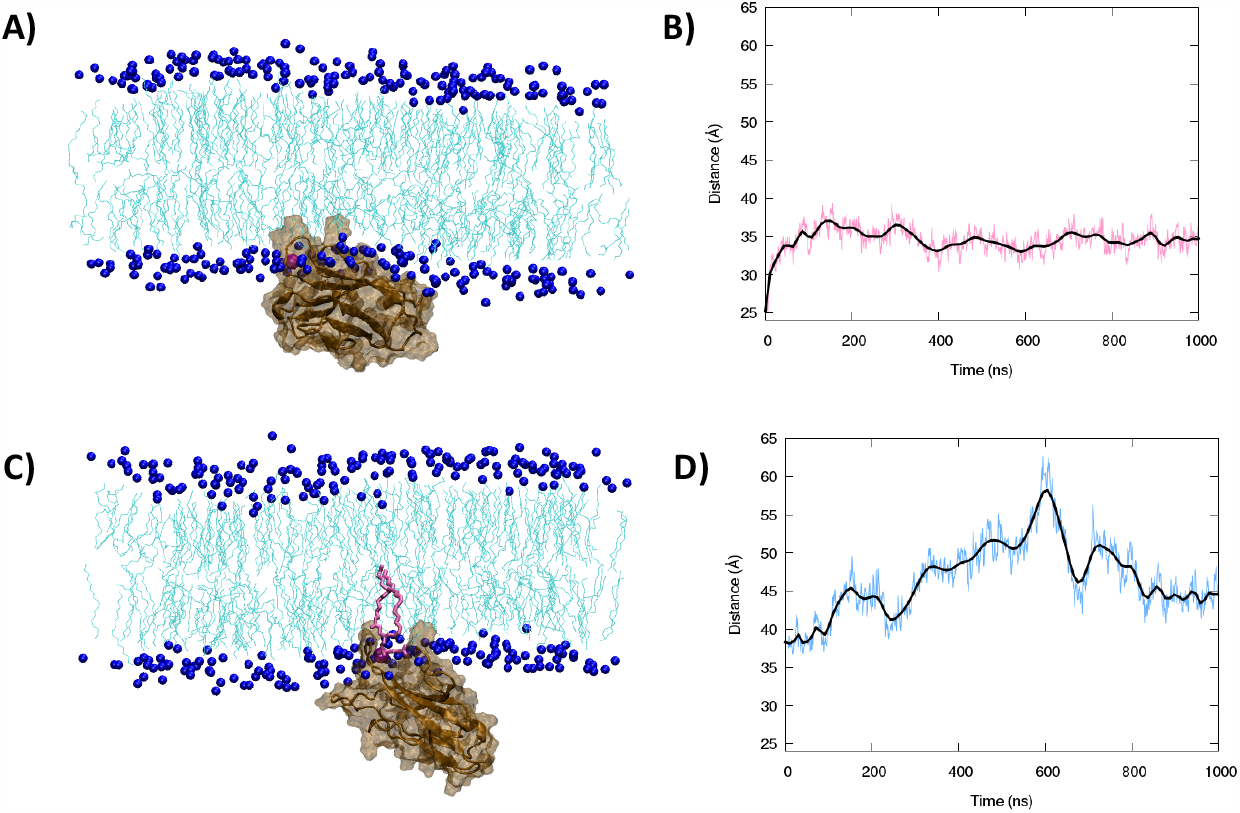
Representative snapshot of the TIM-3 IvG domain interacting with a model lipid membrane for the POPS unbound (A) and bound (B) state. The corresponding time evolution of the distance between the center of mass of the protein and of the lipid bilayer are also given in panel B) and C) for the bound and unbound state, respectively. The protein is represented in cartoon and transparent surface, the position of the lipid polar head is evidenced by plotting the Choline nitrogen atom in van der Waals. The binding pocket is evidenced by the presence of Ca^2+^ ions in purple van der Waals. Finally, in panel C) the bound POPS is represented in licorice.

**Figure 3).**
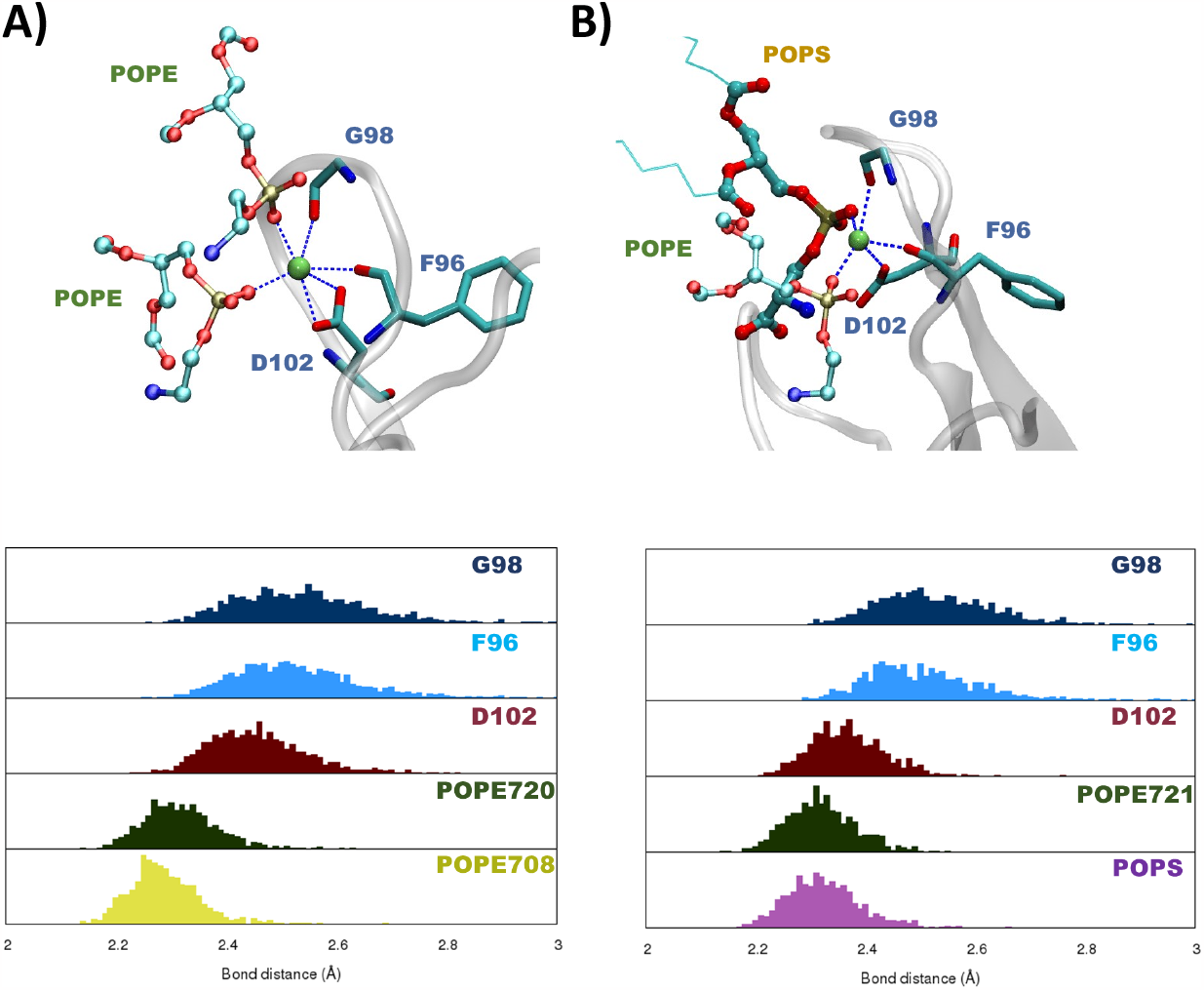
Representative structures evidencing the interactions with the Ca^2+^ ion for the unbound (A) and the bound (B) state. The distribution of the distances with the interacting residues is given in the bottom panel. Note that the presence of POPE lipids interacting with Ca^2+^ without occupying the binding pocket is also highlighted with the former represented in translucid licorice. In case of lipids only the polar heads have been considered to calculate the distance.

While the protein/membrane complex remains persistent upon the binding of POPS polar head some global rearrangements take place. Indeed, a slightly tilting of the IvG domain, with respect to the membrane axis, can be evidenced leading to a straighter orientation of the POPS bound state, compared to the POPS unbound structure, which lies flatter on the membrane hedge. The effects of this different orientation can also be observed on the time evolution of the distance between the center of mass of the protein and of the membrane as reported in Figure 2C, D. Indeed, in the case of the unbound state the distance remains remarkably stable platooning at around 35 Å and experiencing only very limited fluctuations, not exceeding 5 Å. On the other hand, in the POPS bound state the distance is larger and seems to equilibrate at around 45 Å. Even more importantly, rather significant deviations may be evidenced, such as a sharp increase in the distance peaking at around 60 Å at 600 ns. Once again, this sudden increase of the distance is not due to the release of the FG and CC’ loops from the polar head regions, but rather to a conformational and rotational reorientation of the protein relative to the membrane axis. Even if this reorientation can also be related to the long-range signal transduction exerted upon binding, which further triggers the immune system activation, at the moment this hypothesis can be only regarded as speculative, and would necessitate the treatment of the whole TIM-3 instead than the simple IvG domain.

Going from a global to a more local analysis of the IvG behavior, we can clearly see that the Ca^2+^ ion plays a fundamental role in structuring the binding pocket, either in presence or absence of the POPS ligand. In Figure 3 and Table 1 we report the most important residues interacting with the Ca^2+^ ion, and the values of the most crucial distances in both bound and unbound state.

**Table 1).** Average values and standard deviation of the crucial distances involving the Ca2+ in the IvG binding pocket. Note that for lipid only the polar heads have been considered to calculate the distance.

|  | Unbound State | Bound State |
| --- | --- | --- |
| G98:O | 2.5 ± 0.16 | 2.5 ± 0.11 |
| F96:O | 2.5 ± 0.22 | 2.5 ± 0.11 |
| D102:OD1 | 2.1 ± 0.04 | - |
| D102:OD2 | 2.5 ± 0.20 | 2.4 ± 0.06 |
| POPS | - | 2.3 ± 0.06 |
| POPE721 | - | 2.3 ± 0.17 |
| POPE708 | 2.4 ± 0.53 | - |
| POPE720 | 3.8 ± 3.18 | - |

While the Ca^2+^ interaction network appears highly conserved and involve mainly G98, F96 and D102, some differences between the two states may be evidenced. Indeed, while D102 develops two interactions with Ca^2+^ in the unbound state only one of them is conserved upon binding. This is also mostly due to the competition with PdtSer for the occupancy of the binding pocket. Furthermore, as can be appreciated in Figure 3, the interactions with aspartate appears more persistent and the distribution more peaked than those involving glycine and phenylalanine. This aspect is not surprising due to the favorable interactions between negative aminoacids and highly charged cations.

Finally, and interestingly it has to be noted that in the bound state the PdtSer polar head of POPS interacts strongly with Ca^2+^ contributing to its stabilization in the binding cleft. However, in the same state we may also note the presence of a POPE polar head which develops persistent interactions with the divalent cation. The affinity of POPE for Ca^2+^ can be even better assessed in the unbound state, where, because of the absence of POPS, two POPE moieties come in close proximity to the binding pocket and develop highly persistent interactions with Ca^2+^. Yet, as shown in Figure 3A, these two POPE units do not fully occupy the binding cleft, differently from POPS and PdtSer polar head (Figure 2B), thus leaving the site available for the binding with the native ligand. Note that the very high standard deviation of the distance between Ca^2+^ and POPE720 (Table 1) is due to the fact that in the starting conformation the lipid is not in close proximity to the IvG domain, and develops stable interactions only after 300 ns. Interestingly, these results are also coherent with the observation of the crystal structure, with the partial exception of the interaction with N101 which is not conserved.

The interactions determining the insertion of POPS into the binding pocket and its stabilization are, instead, represented pictorially in Figure 4 and quantitatively in Table2, together with their distribution over the span of the MD simulation for the bound state. As already evidenced, the interaction with Ca^2+^, which is highly persistent and presents a rather peaked distribution, appears as one of the leading factors for the binding of the lipid. On the same footing a strong and persistent hydrogen-bond, which is also probably mediated by calcium, takes place between POPS and the backbone nitrogen of D102. Finally, less persistent and more mobile interactions take place with the backbone of Q43 and S42, which occupy the CC’ loop and complete the stabilization of the binding cleft, despite having an average distance greater than 3 Å and a more important standard deviation.

**Table 2).**
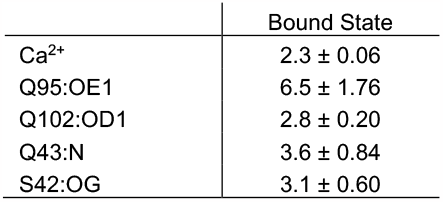
Average values and standard deviation of the crucial distances involving POPS and the binding pocket residues.

|  | Bound State |
| --- | --- |
| $\text{Ca}^{2+}$ | $2.3 \pm 0.06$ |
| Q95:OE1 | $6.5 \pm 1.76$ |
| Q102:OD1 | $2.8 \pm 0.20$ |
| Q43:N | $3.6 \pm 0.84$ |
| S42:OG | $3.1 \pm 0.60$ |

**Figure 4).**
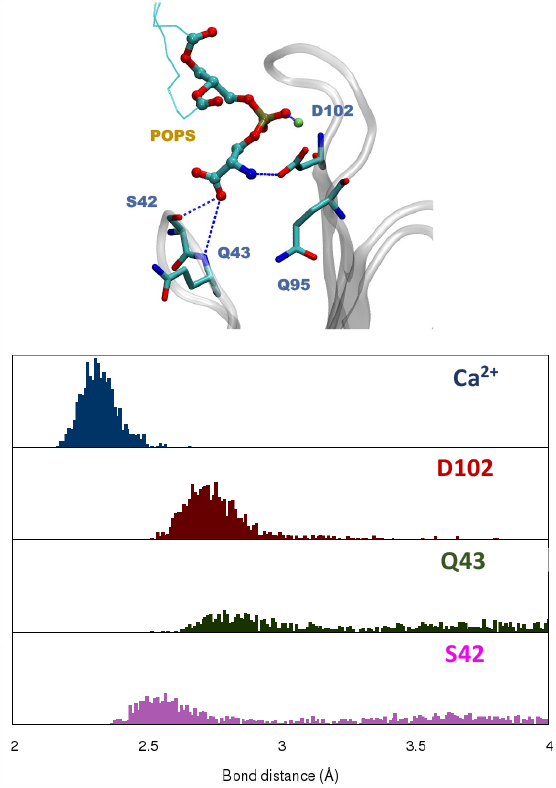
Representations of the main interactions stabilizing the POPS polar head in the binding pocket of the TIM-3 IvG domain. The distributions of the crucial distances are also given in the bottom panel. Note that for POPS only the polar heads have been considered to calculate the distance.

Once again, these results are consistent with the analysis of the crystal structure with the exception of Q95 whose interaction with POPS is not persistent and give raise to a very high standard deviation and an average distance of more than 6.5 Å.

In addition to the results issued from the equilibrium MD simulation in Figure 5 we report the Potential of Mean Force (PMF) giving the free energy profile for the interaction of POPS with the binding cleft of the TIM-3 IvG domain. The PMF is performed by monitoring the distance between the PdtSer polar head and the IvG binding pocket.

**Figure 5).**
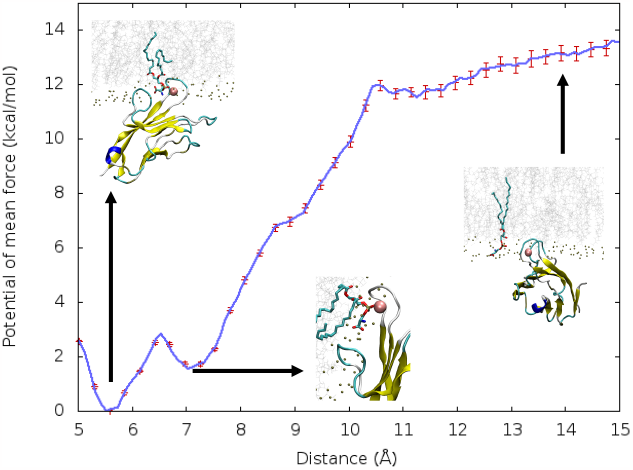
PMF along the collective variable defined as the distance between POPS and the IvG domain binding pocket. Snapshot of the most relevant areas of the PMF are also provided as inlays.

The recognition and the first binding of POPS, resulting on the approach of the lipid to the IvG domain, is almost barrierless with only a small energetic penalty of less than1 kcal/mol appearing at around 10.5 Å. Interestingly, an intermediate state can be observed at 7 Å corresponding to a situation in which POPS is interacting with Ca^2+^, albeit in a still unfavorable orientation to proceed to the further insertion of the polar head in the binding pocket. After overcoming a free energy barrier of about 1.5 kcal/mol PdtSer (and thus POPS) can penetrate deeper in the pocket reaching the absolute minimum conformation. At the minimum of the PMF all the previously described interactions can, indeed, be formed, further confirming the robustness of the equilibrium MD sampling. Importantly, POPS binding results in a free energy gain which amounts to about 12 kcal/mol. Such an important driving force coupled with the almost barrierless profile, points to a very efficient recognition of POPS, which is coherent with the biological role exerted by TIM-3.

The binding of POPS, and PdtSer, may also affect the organization of the IvG domain and, in particular, the distance between the constitutive loops of the domain. The time series of the distances between the loops (see SI for the precise definition) are reported in Figure 6. We may see that the bound state presents a significant decrease of its flexibility, as evidenced by the reduced fluctuations in the loop distances. Unsurprisingly, the FC loop which largely encompasses the binding pocket appears as the most affected. The most striking difference is visible for the FG-BC distance which is strongly stabilized at around 20 Å in the bound state, while it exhibts large deviations and fluctuations in the unbound system, spanning the 14-22 Å range. Interestingly, the evolution of the loop flexibility has also been pointed as one of the structural reasons behind TIM-3 activation, and hence its biological role.^[44]^

**Figure 6).**
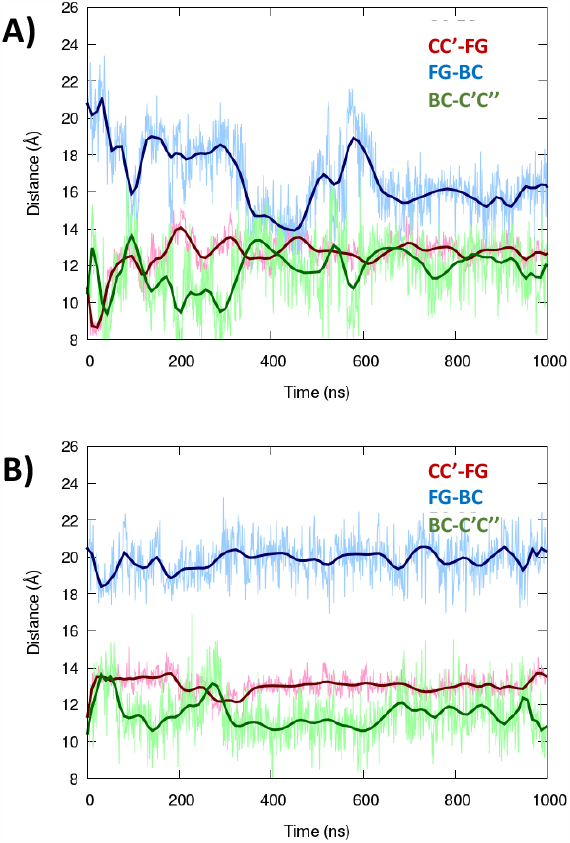
Time evolution of the distances between the IvG loops as defined in SI, for the unbound (A) and bound (B) state.

Finally, since most of the experimental structure of the bound IvG only involves the lipid polar head (PdtSer) without the hydrophobic tails, it is important to assess the conformational flexibility of the bound lipid core and its eventual time evolution.

As seen in Figure 7, the lipid tails remain largely oriented parallel to the main membrane axis, therefore they minimize possible interactions with the loops, and hence eventual steric clashes. Yet, the presence of the lipid tails seems to partially affecting the positioning of the polar head in the binding pocket, which appears less profoundly inserted than in the crystal structure.

**Figure 7).**
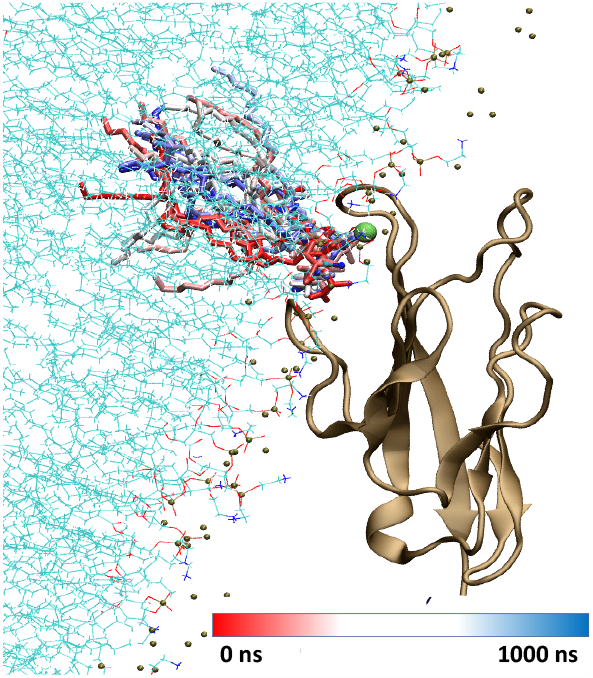
Structure of the IvG domain interacting with the lipid bilayer. The time evolution of the position of the bound POPS lipid tails is shown (evolution from blue to red).

## Conclusion

The interaction between the IvG domain of TIM-3 with a model lipid membrane has been characterized at a molecular level thanks to the use of all-atom MD simulations exceeding the μs time-scale. We have shown that the IvG domain forms stable peripheral complexes with the lipid bilayer, however the interaction is loose enough to permit the sliding of the protein domain over the membrane plane. This specificity may help encountering the natural TIM-3 substrate, i.e. a PdtSer containing lipid. In our case we have shown that the interaction with a PdtSer containing POPS lipid is highly favorable and proceeds via the important participation of the Ca^2+^ ion in the IvG binding pocket, while being assisted by the formation of an extended hydrogen bond network, involving also the protein backbone. Notably, the role of Ca^2+^ is also extremely important in shaping the binding pocket and its rigidity. The use of enhanced sampling MD has also shown that the interaction of POPS with the IvG domain, leading to the insertion of PdtSer in its binding pocket, is favorable and has a thermodynamic driving force of about 12 kcal/mol. Interestingly, the insertion is almost barrierless, even if a very labile intermediate can be observed, corresponding to the first interaction between POPS and the Ca^2+^ prior to a favorable reorientation.

The interaction of POPS is also correlated to some conformational variation of IvG, both global and local. Indeed, some titling between the protein and the membrane axis, ultimately resulting in an increase of the distance between the two centers of mass, have been evidenced by our simulations. Furthermore, a rigidification of the IvG loops upon the binding is observed, while the lipid tails of the bound lipid remain parallelly to the membrane axis. This aspect can be interesting for the modulation of TIM-3 activity by modified lipids bearing photoswitching moieties. Indeed, as already shown experimentally,^[49]^ the E/Z isomerization may induce conformations of the lipids which will decrease the affinity due to partial clashes with the IvG domain. This topic will be the subject of a future contribution, and will strongly profit from the knowledge of the molecular bases driving the interaction as described in this contribution.

Overall, we have provided the first comprehensive elucidation of the molecular level functioning of a key actor of the immune system, which is present in T-cell, and whose deregulation in cancer cells suggests its strong role in shaping tumor microenvironment,^[40,41]^ thus constituting a key target for possible immunotherapy.^[32]^

## Computational Methodology

The CHARMM-GUI utility^[50,51]^ has been used to generate an initial lipid bilayer mimicking the composition of eukaryotic cells, interacting with TIM-3 IvG domain. The chosen lipids have been POPC, POPE, POPS and cholesterol, with respective molar ratios of 30%, 25%, 15% and 30%, a total of 200 lipids were placed in each leaflet. In addition, a water buffer of 30 Å was also introduced including a 0.15 M physiological concentration of K^+^ and Cl^-^ ions. The structure of the IvG domain of the murine Tim-3 complexed with a PdtSer polar head has been retrieved from the PDB database (PDB: 3KAA.^[29]^ and used as the starting conformation. Notably, the murine model shows 76% homology with the human analogous. The PtdSer polar head present in the crystal structure has been removed, while maintaining the crystal Ca^2+^. The protein has been placed in the water bulk in proximity to the polar heads to build the initial system. All the classical MD simulations have been performed using NAMD^[52,53]^ and subsequently analysed and visualised with VMD.^[54]^

The lipids have been represented using the Amber “Lipid14” force field,^[55]^ while water has been described with TIP3P^[56,57]^ and the IvG domain of TIM-3 by the Amber ff14SB^[58]^ protein force field. In addition, hydrogen mass repartitions^[59]^ has been consistently used, allowing, in combination with Rattle and Shake algorithms,^[60]^ the use of a 4.0 fs time-step for the integration of the Newton equations of motion. Before production, all the systems underwent a minimization process, followed by thermalization and equilibration. This involved the gradual removal of positional constraints on non-water heavy atoms over a total span of 12 ns. We maintained a constant temperature of 300 K (NVT ensemble) throughout both equilibration and production MD simulations, ensuring the lipid membrane remained in its liquid phase. The equilibration was conducted in the isothermal and isobaric (NPT) ensemble to allow density adjustments. Temperature and pressure conservation were enforced using the Langevin thermostat^[61]^ and barostat,^[62]^ respectively. The production run for this system, representing the POPS-unbound state, was propagated for a total simulation time of 1 μs.

From a representative snapshot of the unbound state (extracted after 218 ns) a second system was built by performing steered MD (SMD),^[63]^ to enforce the penetration of a POPS lipid into the FG-CC’ binding pocket. This was performed by applying a harmonic force of 10 kcal/mol over 200 000 steps with timestep of 1 fs, while the collective variable was chosen as the distance between the centre of mass of the closest POPS polar head (PdtSer) and of the reactive pocket. Subsequently to the SMD procedure, we performed an unconstrained equilibrium MD simulation of 1 μs, ensuring that a proper reorganization and equilibration may take place, thus leading to the bound-POPS equilibrated state. For both bound and unbound states the stability was also estimated by the analysis of the root-mean-square deviation (RMSD)^21^ for both the protein and the lipid bilayer.

Finally, Umbrella Sampling (US)^[64]^ has been used to obtain the potential of mean force (PMF) associated with the binding of POPS, via its PdtSer polar head, and the binding pocket in the FG-CC’ loop of IvG. To this aim we stepwise enlarged the distance between POPS and the protein, starting from the bound state. The collective variable chosen for US was the same as for SMD and involved the distance between the centre of mass of the bound POPS polar head (PdtSer), and the binding pocket, i.e. the Ca^2+^ ion as well as the S42, Q43, Q95, and D102 residues. A graphical representation of the collective variable is provided in Supplementary Information (SI). The collective variable was increased from 5.0 to 15.0 Å in independent windows separated by 0.5 Å, each window was propagated for either 15 or 30 ns, to assure a good equilibration. The final PMF profile was obtained combining the different windows with the Weighted Histogram Analysis Method (WHAM).^[65]^ As shown in Supplementary Information a good overlap between windows necessary for the WHAM reconstruction is assured.

## Supporting Information

Plot of the RMSD time evolution for the bound and unbound state, definition of the collective variable, and of the loop distance descriptor.

## Acknowledgements

The authors thank GENCI and Explor computing centers and the Platform P3MB for computational resources. A.M. thanks ANR and CGI for their financial support of this work through Labex SEAM ANR 11 LABX 086, ANR 11 IDEX 05 02. The support of the IdEx “Université Paris 2019” ANR-18-IDEX-0001. Support from the Regional Development Fund of the European Union (Programme opérationnel FEDER-FSE Lorraine et Massif des Vosges 2014-2020/ “Fire Light” project: Photo-bio-active molecules and nanoparticles”) for financial support.

## Entry for the Table of Contents

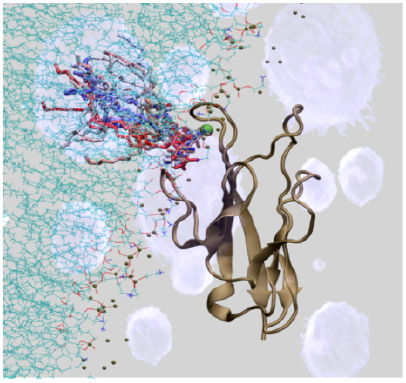

We report molecular modeling and simulation of the specific recognition of Phosphatidyl serine-containing lipids and TIM-3 immune system agent, which is particularly present in natural killer and T-cells. We also identify the thermodynamic of the interaction process, using enhanced sampling molecular dynamics.

Institute and/or researcher Twitter usernames: @antoniomonari, @ITODYS_lab, @L2CM_UMR7053, @univ_paris_cite

## Supplementary Information for

**Figure S1).**
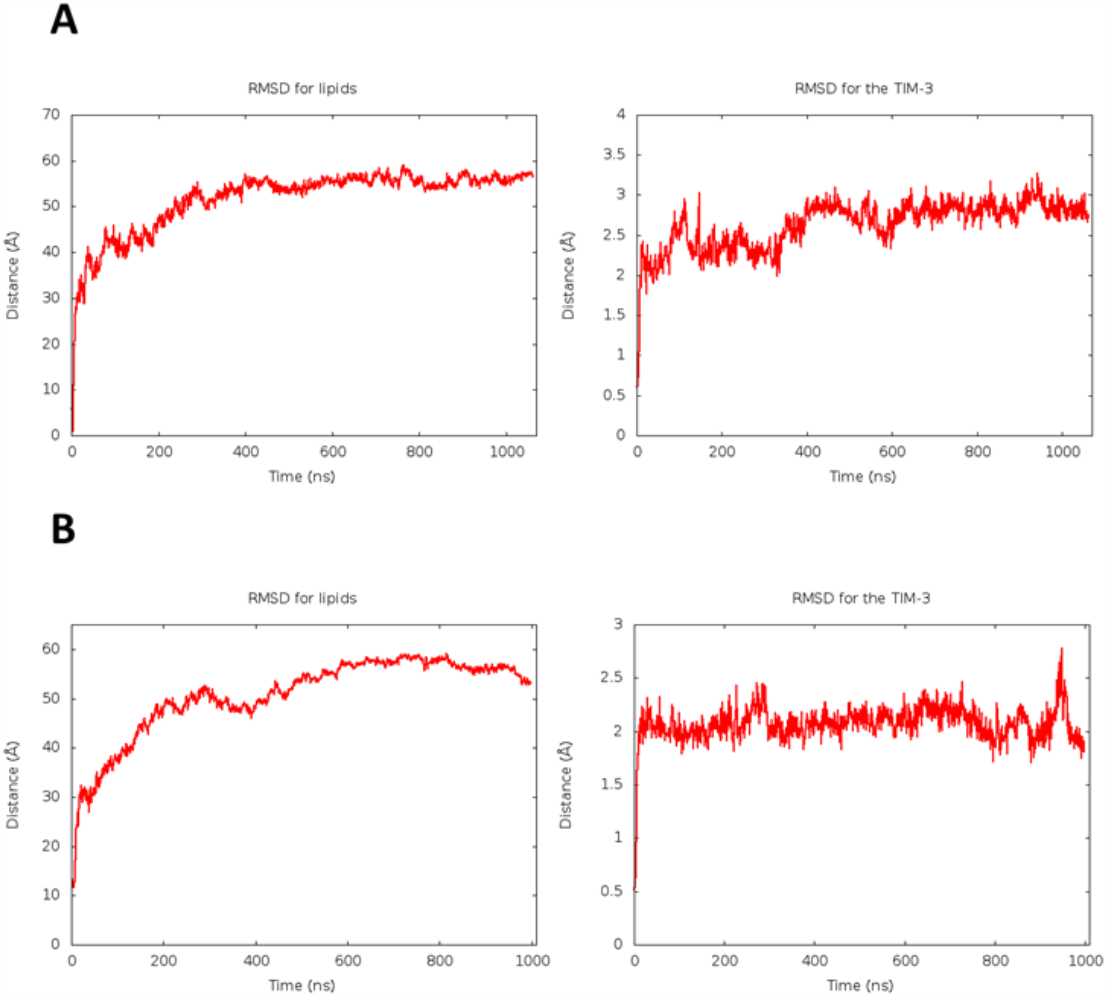
Time evolution of the root mean square deviation (RMSD) for POPS, POPE, POPC, cholesterol lipids and TIM-3’s IgV domain calculated for the unbound (A) and bound (B) state, on the 1 μs MD trajectory.

**Figure S2).**
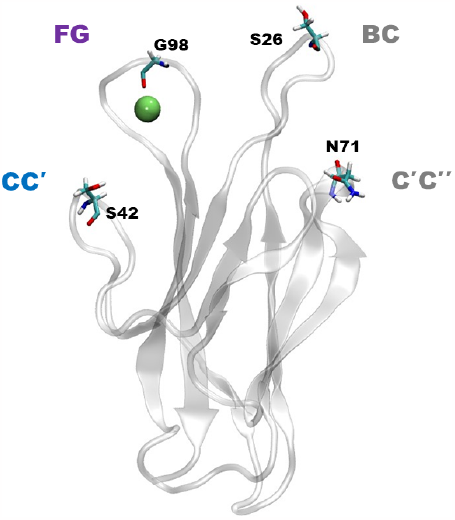
Snapshots showing the protein residue whose distance is used to define the global loop distance. Namely S42—G98 for loops CC’—FG, G98—S26 for FG—BC, and S26—N71 fro the BC—C’C’’ loop.

**Figure S3).**
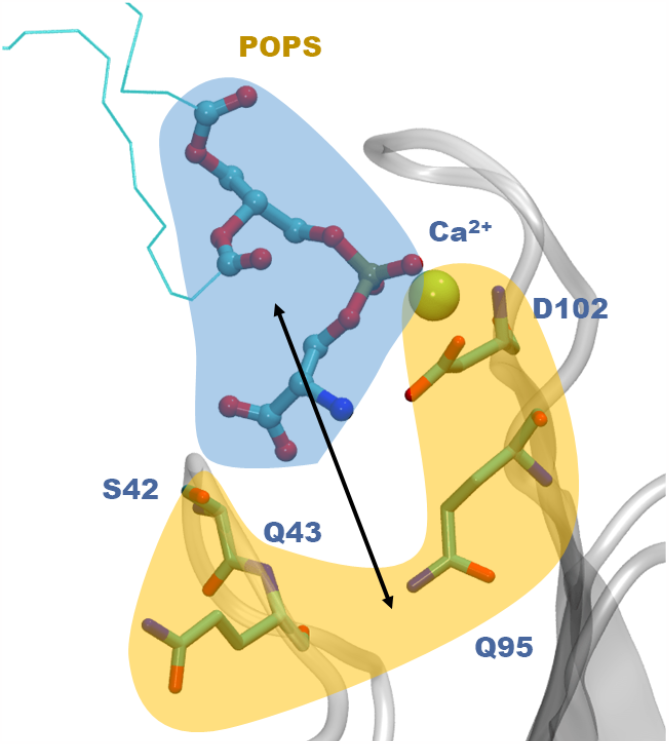
Pictorial representation of the collective variable used for SMD and US simulations. The first group of atoms comprises the polar head of POPS (blue shade) and the second one the Ca^2+^ ion and the S42, Q43, Q95, D102 residues. The distance between of the center of mass of the two groups has then be taken into account.

**Figure S4).**
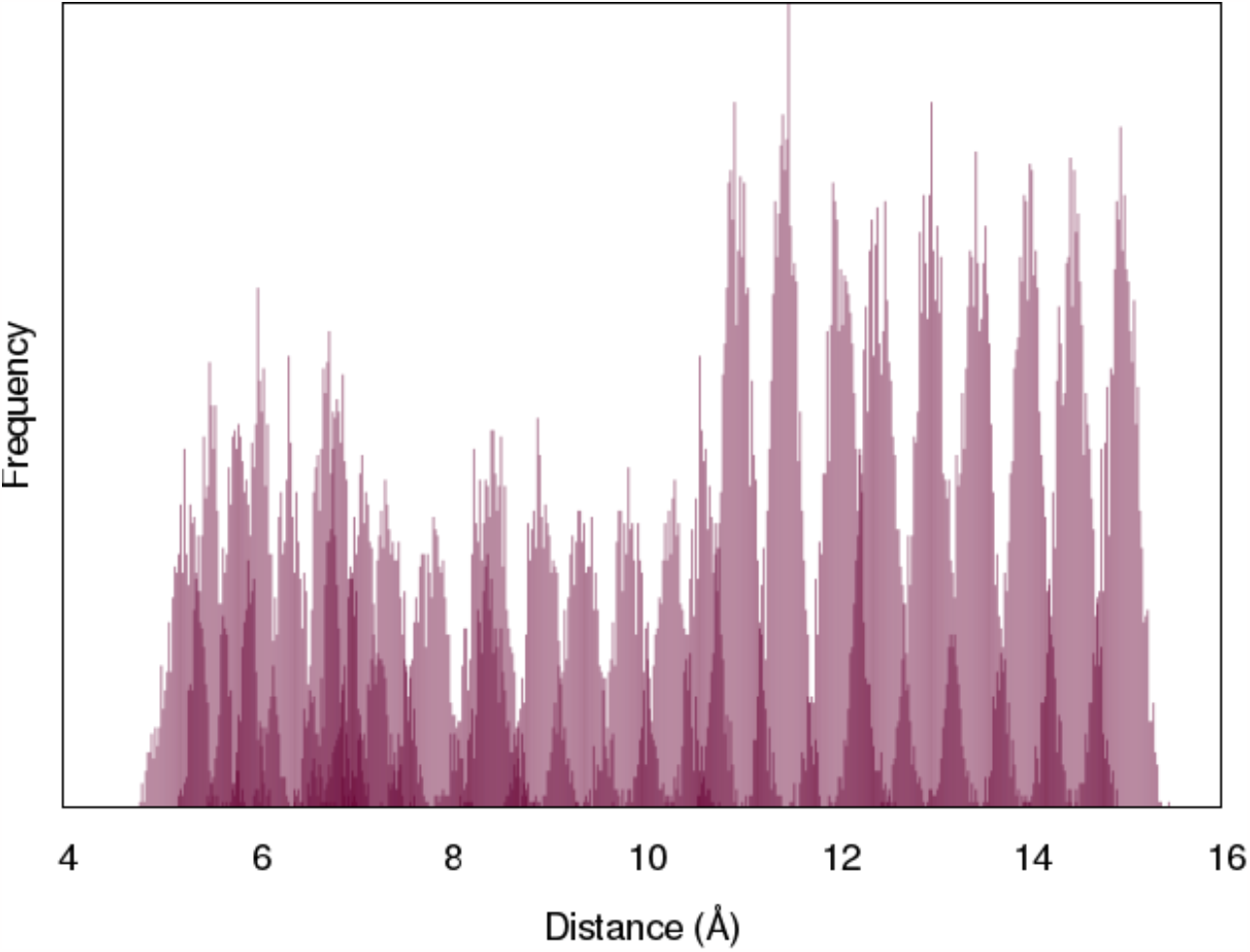
Histogram showing the distribution of the collective variable values among the different windows used for the US.

